# Using genetically encoded heme sensors to probe the mechanisms of heme uptake and homeostasis in *Candida albicans*

**DOI:** 10.1101/2020.07.26.221606

**Authors:** Ziva Weissman, Mariel Pinsky, Rebecca K. Donegan, Amit R. Reddi, Daniel Kornitzer

## Abstract

*Candida albicans* is a major fungal pathogen that can utilize hemin and hemoglobin as iron sources in the iron-scarce host environment. While *C. albicans* is a heme prototroph, we show here that it can also efficiently utilize external heme as a cellular heme source. Using genetically encoded ratiometric fluorescent heme sensors, we show that heme extracted from hemoglobin and free hemin enter the cells with different kinetics. Heme supplied as hemoglobin is taken up via the CFEM (Common in Fungal Extracellular Membrane) hemophore cascade, and reaches the cytoplasm over several hours, whereas entry of free hemin via CFEM-dependent and independent pathways is much faster, less than an hour. To prevent an influx of extracellular heme from reaching toxic levels in the cytoplasm, the cells deploy Hmx1, a heme oxygenase. Hmx1 was previously suggested to be involved in utilization of hemoglobin and hemin as iron sources, but we find that it is primarily required to prevent heme toxicity. Taken together, the combination of novel heme sensors with genetic analysis revealed new details of the fungal mechanisms of heme import and homeostasis, necessary to balance the uses of heme as essential cofactor and potential iron source against its toxicity.

## INTRODUCTION

*Candida albicans* is a commensal of human mucosal surfaces, as well as an opportunistic pathogen that can cause superficial mucosal infections among immunocompetent individuals and lethal systemic infections among immunocompromised patients (van de Veerdonk *et al*., 2010; Jabra-Rizk *et al*., 2016). It is the most prevalent systemic fungal pathogen, responsible yearly for close to a million cases with high (∼40%) mortality (Kullberg and Arendrup, 2015; Bongomin *et al*., 2017). Like all commensal and parasitic microorganisms, *C. albicans* must be able to extract essential nutrients from the host environment including iron, for nourishment (Ramanan and Wang, 2000). Host iron is sequestered in iron-binding proteins and is therefore normally unavailable for parasitic microorganisms and furthermore, the host actively withholds iron in blood and tissues during inflammation, to further restrict pathogen proliferation (Weinberg, 1975; Cassat and Skaar, 2013; Ganz and Nemeth, 2015). Therefore, in order to acquire iron, successful pathogens have evolved mechanisms to extract it from host proteins. Heme (iron protoporphyrin IX) represents greater than 70% of the human host’s iron quota due to its role as a cofactor for the abundant oxygen transport blood protein, hemoglobin (Hb) (Ganz and Nemeth, 2015).

Consequently, pathogens often target hemoglobin as an iron source in the iron-poor host environment (Wandersman and Delepelaire, 2004; Caza and Kronstad, 2013; Choby and Skaar, 2016; Bairwa *et al*., 2017; Hare, 2017; Roy and Kornitzer, 2019). Notably, the ability to utilize heme as iron source is not restricted to pathogens: the widespread occurence of heme in live and dead organic matter makes it a potential source of iron in various environments (Contreras *et al*., 2014; Roy and Kornitzer, 2019).

*C. albicans* is able to utilize external heme, both in the form of free hemin (the chloride ion of Fe^3+^ heme), as hemoglobin heme, or as heme complexed with albumin, as iron sources (Moors *et al*., 1992; Pinsky *et al*., 2020). Hemoglobin and albumin-complexed heme utilization requires a cascade of extracellular hemophores, both soluble and cell wall- and plasma membrane-anchored, that bind heme via a CFEM (Common in Fungal Extracellular Membrane) domain (Nasser *et al*., 2016). The CFEM hemophores Csa2, Rbt5 and Pga7 capture the heme and transfer it across the cell envelope to the endocytic pathway (Weissman and Kornitzer, 2004; Kuznets *et al*., 2014; Pinsky *et al*., 2020) (reviewed in (Roy and Kornitzer, 2019)). This pathway also requires parts of the ESCRT (Endosomal Sorting Complexes Required for Transport) system, an endocytosis-specific myosin, and the vacuolar ATPase (Weissman *et al*., 2008). Free hemin utilization at low concentration similarly requires the CFEM hemophore cascade, but at higher hemin concentration, the CFEM system can be bypassed (Weissman and Kornitzer, 2004; Kuznets *et al*., 2014; Pinsky *et al*., 2020).

In order for the heme-iron to be utilized, the iron atom must be extracted from protoporphyrin IX. Heme oxygenase enzymes, which are widely conserved among eukaryotes, catalyze the O_2_ dependent degradation of heme, releasing CO, iron, and biliverdin, and play an essential role in the recycling of cellular iron (Maines, 1988; Poss and Tonegawa, 1997; Ponka, 1999). In *C. albicans*, a heme oxygenase homolog, Hmx1, was identified and was found to be involved in utilization of free hemin as an iron source (Santos *et al*., 2003; Pendrak *et al*., 2004).

Aside from its role as a potential source of nutritional iron, heme also serves as an essential cofactor and signaling molecule (Ponka, 1999; Mense and Zhang, 2006; Ortiz de Montellano and Begley, 2007). However, the same characteristics that make heme a potent cofactor, including its lewis acidity, redox activity and hydrophobicity, also cause heme to be toxic. This toxicity is in part due to the misincorporation of heme into proteins and membranes and deleterious redox reactions catalyzed by heme (Gutteridge and Smith, 1988; Solar *et al*., 1991). In order to ensure heme is bioavailable for heme dependent processes, while also mitigating its cytotoxicity, labile heme (LH) is expected to be tightly regulated. In addition to their role in iron recycling, mammalian heme oxygenases are also thought to be involved in preventing heme toxicity if cytosolic heme levels reach dangerously high levels (Maines, 1988; Ponka, 1999). However, the molecules and mechanisms that regulate LH in *C. albicans*, and generally any organism, are not well understood. Moreover, the contribution of exogenously scavenged heme towards LH pools and its bioavailability for heme dependent processes, *i*.*e*. whether *C. albicans* can utilize scavenged heme as source of cellular heme, was not known.

We show here that exogenously scavenged heme can in fact serve as a heme source in *C. albicans* and contribute to the LH pool. Ratiometric fluorescent sensors for LH were recently developed and utilized in *Saccharomyces cerevisiae, E. coli*, and mammalian cell lines to probe heme trafficking and dynamics (Hanna *et al*., 2016; Hanna *et al*., 2017; Hanna *et al*., 2018; Sweeny *et al*., 2018; Martinez-Guzman *et al*., 2020). Adapting these heme sensors to *C. albicans*, we measured the effects of iron starvation, heme depletion, and extracellular heme supplementation on intracellular heme levels. We find that cytoplasmic LH can be significantly affected by these factors, particularly in the absence of the heme oxygenase Hmx1.

## RESULTS

### Hemin and hemoglobin can serve as heme sources for *C. albicans*

In order to test whether hemin and hemoglobin can also serve as heme sources, in addition to iron sources, for *C. albicans*, we inhibited heme synthesis in this organism by interfering with the first step of the endogenous heme biosynthetic pathway, which is carried out by δ-aminolevulinic acid (ALA) synthase, Hem1. Inhibition of heme synthesis was achieved by mutation or down-regulation of *HEM1*. Although heme is an essential cofactor, a *hem1*^*-/-*^ strain can be kept alive by supplementing the growth media with ALA, the Hem1 reaction product. Alternatively, heme biosynthesis was inhibited using a strain where one *HEM1* allele is deleted and the other is under the regulatable TET-off promoter (Roemer *et al*., 2003).

As shown in Fig. 1A, shutting off the *HEM1* gene using the TET-off promoter by addition of doxycycline does prevent growth of colonies, indicating that *HEM1* expression is below that required for cell growth under these conditions. Provision of hemin in the medium led to growth recovery, but no growth was seen with hemoglobin as a heme source. We wondered whether the lack of growth on hemoglobin could be due to a lack of induction of the heme-uptake pathway. Indeed, the known genes specifically required for extraction of heme from hemoglobin and heme uptake, the CFEM hemophore genes *RBT5, PGA7* and *CSA2*, are among the highest-induced genes under iron starvation conditions (Lan *et al*., 2004; Weissman and Kornitzer, 2004; Chen *et al*., 2011; Roy and Kornitzer, 2019). We thus also tested growth on hemin and hemoglobin in medium supplemented with 1 mM ferrozine, an iron chelator that imposes partial iron starvation on the cells and causes induction of the heme uptake genes (Weissman and Kornitzer, 2004). In the presence of ferrozine, growth on hemoglobin as an external heme source was much improved (Fig. 1A). The ferrozine requirement was also seen when the cells were grown in liquid media with increasing hemin or hemoglobin concentrations. Hemin and hemoglobin allowed growth of the *hem1*-strain above 1.25 µM and 0.16 µM, respectively, in iron starvation medium, but a higher concentration of hemin was needed in regular medium, and no growth was recorded on hemoglobin in regular medium (Fig. 1B).

**FIG 1.**
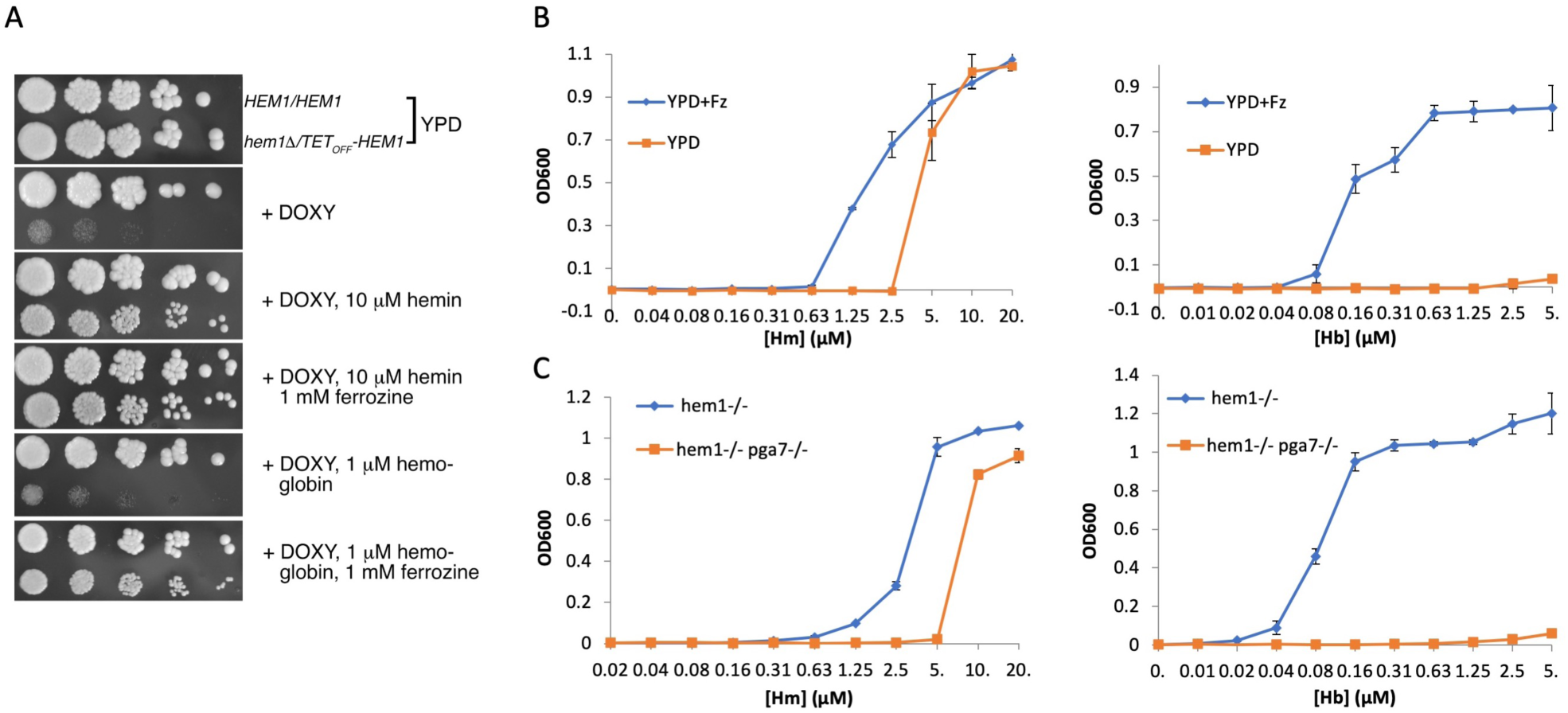
*C. albicans* can efficiently utilize external hemin as cellular heme source. A. Dilutions of a wild-type strain (CaSS1) and a *hem1*^*-*^*/TEToff-HEM1* strain were spotted on the indicated plates without or with 50 µg/ml doxycycline (DOXY), and the indicated hemin or hemoglobin supplements. The plates were incubated for 2 days at 30°C. B. The *hem1*^*-*^ */TEToff-HEM1* strain was grown in YPD medium with 50 µg/ml doxycycline, with or without 1 mM ferrozine as indicated, and with the indicated concentrations of hemin or hemoglobin. The cultures were grown at 30°C for 3 days. Each data point represent the average of three independent cultures, and the error bars represent the standard deviation. C. The *hem1*^*-/-*^ (KC1088) and *hem1*^*-/-*^ *pga7*^*-/-*^ (KC1111) strains were grown in YPD medium with 1 mM ferrozine, and with the indicated concentrations of hemin or hemoglobin. The cultures were grown and measured like in B.

A *hem1*^*-/-*^ strain was next constructed by CRISPR mutagenesis. The *hem1*^*-/-*^ strain was unable to grow in the absence of ALA, as expected, but growth of this strain was restored by addition of hemin or hemoglobin (Fig. 1C). We then tested whether utilization of external heme as a heme source required the same hemophore system as heme-iron utilization by deleting *PGA7*, a deletion which causes a strong heme-iron utilization defect (Kuznets *et al*., 2014). As seen in Fig. 1C, in the absence of *PGA7*, hemoglobin utilization as a heme source was abolished, while hemin utilization was reduced, similar to what was previously observed for hemin utilization as an iron source in this mutant (Kuznets *et al*., 2014; Pinsky *et al*., 2020). In addition, we previously found that addition of albumin facilitates heme-iron utilization (Pinsky *et al*., 2020). We find that albumin similarly promotes utilization of heme as a heme source (Fig. S1).

### The heme oxygenase Hmx1 is not essential for heme-iron utilization

Since *C. albicans* can take up external heme and use it as a heme source, we next addressed the specific requirements for utilization of heme as an iron source. Most eukaryotic cells possess a heme-degrading heme oxygenase enzyme to protect themselves against the toxicity of excess heme. Previous work had shown that *HMX1* is required for utilization of hemin as an iron source in *C. albicans* (Santos *et al*., 2003). We thus tested whether hemoglobin-iron utilization also requires Hmx1. We found however that the utilization of both hemin and hemoglobin as iron sources are only weakly dependent on Hmx1. Using a published strain set (Navarathna and Roberts, 2010), when tested on plates, a small growth defect on either hemoglobin or hemin was detected in the *hmx1*^*-/-*^ strain, compared to the starting strain or the complemented *HMX1* strain (Fig. 2A). A similar small growth defect was detected using another strain set (Fig. S2). In liquid, no significant growth difference could be detected with either hemin or hemoglobin as iron sources (Fig. 2B). We thus conclude that Hmx1 is not essential for heme-iron utilization in *C. albicans*.

**FIG 2.**
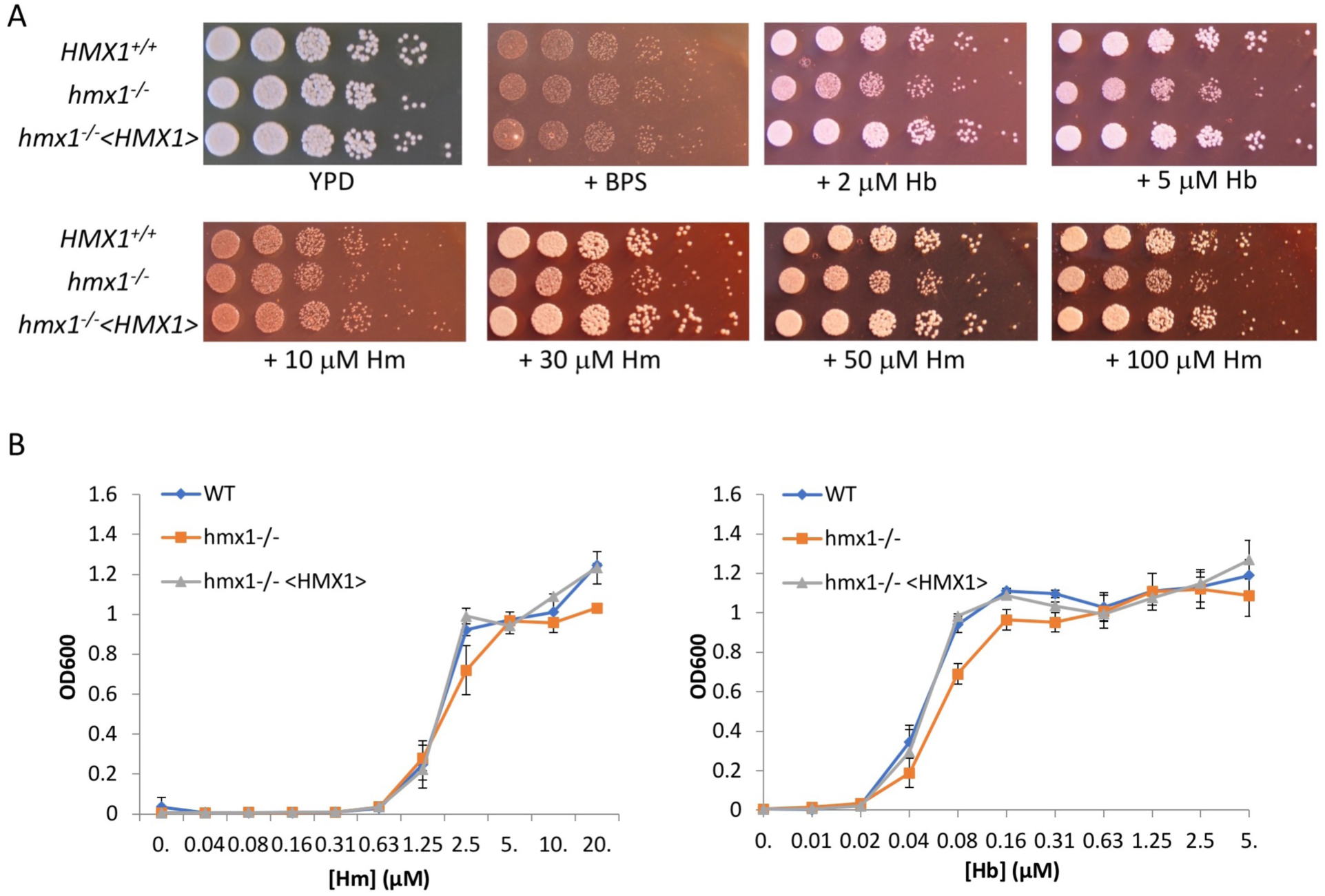
Cells lacking the heme oxygenase Hmx1 are still able to utilize hemin and hemoglobin as iron sources. A. Dilutions of a wild-type strain (KC1065), of a *hmx1*^*-/-*^ mutant (KC1066), and of a *hmx1*^*-/-*^ <*HMX1*> reintegrant (KC1067) were spotted on YPD or on YPD + 1 mM BPS, without or with the additions of the indicated hemoglobin and hemin concentrations, and incubated at 30°C for 2 days. B. The same set of strains was tested for growth in liquid with hemin or hemoglobin as iron sources. Cultures were grown at 30°C for 2 days.

### Monitoring cytoplasmic labile heme concentration

The data presented above show that *C. albicans* is an organism that can either synthesize its own heme or utilize hemoglobin and hemin as heme sources. Such an organism must carefully balance *de novo* heme synthesis and heme uptake in order to prevent accumulation of toxic levels of intracellular heme. In order to measure cytoplasmic LH levels under different regimes of internal and external heme supply, we adapted a set of fluorescent ratiometric cellular heme probes (Hanna *et al*., 2016; Hanna *et al*., 2017) to *C. albicans*. The cellular heme probe consists of two fluorescent domains, an mKATE2 red-fluorescing domain and a green fluorescent protein (eGFP) domain (Fig. S3A). Adjacent to the eGFP domain is a cytochrome domain that can bind heme. Binding of heme strongly quenches the eGFP fluorescence but minimally affects mKATE2 fluorescence. The ratio of eGFP to mKATE2 fluorescence gives a measure of the extent of heme binding to the probe, which in turn reflects the concentration of cytoplasmic LH. To extend the dynamic range of detection, three heme sensors are used. The high affinity sensor, wild-type HS1, binds heme using Met and His axial ligands of the cytochrome domain and exhibits dissociation constants of 3 nM for ferric heme (*K*_D_^Fe(III)^) and ∼1 pM for ferrous heme (*K*_D_^Fe(II)^) (Hanna *et al*., 2016; Hanna *et al*., 2017). The moderate affinity heme sensor, HS1-M7A, has the axial methionine ligand mutated to alanine and exhibits weaker heme affinities; *K*_D_^Fe(III)^ ∼ 1 μM and *K*_D_^Fe(II)^ = 25 nM. HS1-M7A,H102A, a double mutant (DM) which has both the methionine and histidine axial ligands mutated to alanine, cannot bind heme and serves as a control for determining heme-independent changes to sensor fluorescence ratios. As shown in Fig. S3B, wild-type HS1 expressed in *C. albicans* exhibits an eGFP/mKATE2 sensor ratio of ∼2.7, which is distinctly lower than the sensor ratio of the non-heme binding variant HS1-M7A,H102A (DM), ∼17.3. On the other hand, HS1-M7A exhibits a similar sensor ratio to HS1-M7A,H102A. These results were confirmed by single cell analysis using fluorescence microscopy (Fig. S3C). Together, these data suggest that HS1 has a sufficiently high binding affinity to bind and sense heme *in vivo*, whereas HS1-M7A binds heme too weakly to monitor heme in cells under normal growth conditions.

By adapting previously established sensor calibration procedures to *C. albicans* (see Materials and Methods), we find that the wild-type HS1 sensor is ∼ 80% loaded with heme, while the HS1-M7A sensor is < 10 % heme loaded in synthetic complete (SC) medium. In principle, the fractional heme occupancy of HS1 can be used to estimate the concentration of buffered LH concentrations (Hanna *et al*., 2016; Hanna *et al*., 2017). However, due to the uncertainty in the oxidation state of LH and the differences in affinity of HS1 for reduced vs. oxidized heme, the concentration of LH cannot be accurately determined. Based on the heme binding affinities of HS1 and its fractional heme occupancy, we estimate [LH] being between 5 pM – 16 nM, with the lower and upper bounds defined by assuming that LH is either 100% reduced or oxidized, respectively. Given these limitations, we use fractional heme occupancy as a measure of relative cytoplasmic LH concentration under different conditions rather than absolute concentrations.

### Kinetics of heme influx

In a first set of experiments, we measured the heme sensor fluorescence ratios in cells grown in regular medium vs. partial iron limitation by addition of the iron chelator ferrozine. Since heme binding quenches eGFP fluorescence, maximal eGFP/mKATE2 fluorescence ratio indicates low heme, and a decrease in ratio indicates an increased cytoplasmic free heme concentration. As seen in Fig. 3A, in the absence of any external hemin or hemoglobin, wild-type cells grown in regular medium show high heme occupancy (∼ 90%) of the wild-type (WT) HS1 sensor, but low heme occupancy (∼ 17%) of the HS1-M7A sensor. In cells grown in the presence of ferrozine, wild-type HS1 occupancy is lower (∼ 30%), indicating that iron-starved cells have a lower LH concentration in the cytoplasm. Addition of increasing concentrations of hemin led to increasing WT HS1 heme occupancy at 5 and 10 µM, and at 10 µM, the lower affinity HS1-M7A sensor became partially occupied with heme as well. In contrast, 10 µM hemoglobin (= 40 µM heme equivalents, since hemoglobin is tetrameric) did not affect intracellular LH concentration in regular medium. In iron starvation medium however, hemin addition increased WT HS1 heme occupancy with 2.5, 5 and 10 µM hemin, and the HS1-M7A sensor also became significantly heme-loaded in 10 µM hemin. With hemoglobin, in iron starvation medium, in contrast to the situation in regular medium, 10 µM hemoglobin led to an increase in WT HS1 heme occupancy. These observations are consistent with the conclusions of the experiments measuring restoration of growth of the *hem1*^*-/-*^ mutant, namely that hemoglobin-heme can only enter the cells when the CFEM heme-uptake pathway is induced by iron starvation, whereas hemin can also penetrate cells in the absence of iron starvation, albeit less efficiently.

**FIG. 3.**
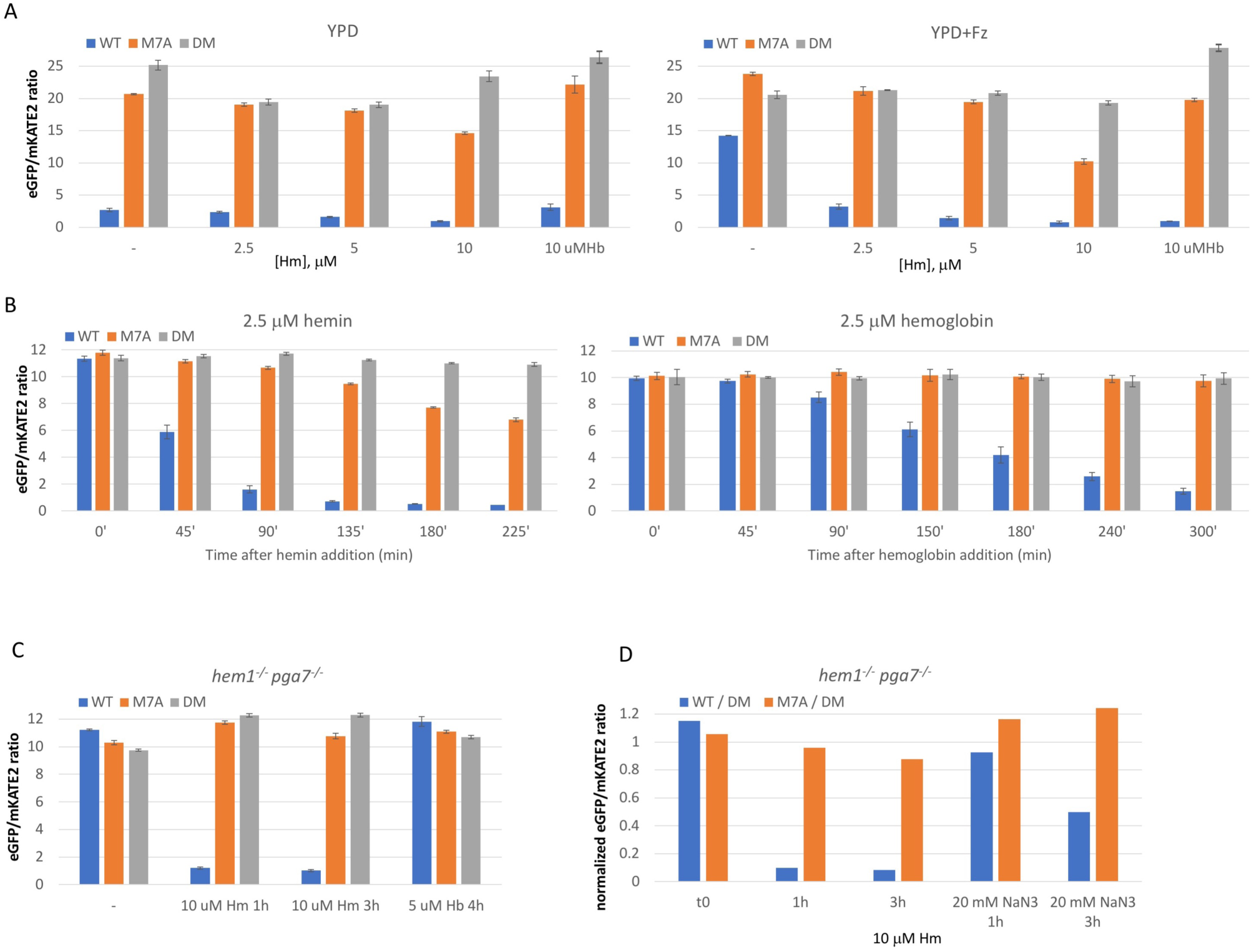
Effect of growth medium and exposure to hemin and hemoglobin on intracellular free heme concentrations. A. Cells (KC1262) expressing the wild-type HS1 heme sensor (WT), the M7A cytochrome mutant, or the M7A H102A cytochrome double mutant (DM), were grown overnight in YPD with or without 1 mM ferrozine, then diluted in the same media supplemented with the indicated amounts of hemin or hemoglobin, and grown for another 5 hours before measuring the fluorescence ratios. Each data point is the average of three independent cultures, two samples each. The value measured is the heme-quenchable eGFP fluorescence divided by the non-quenchable mKATE2 fluorescence. Error bars indicate the standard deviation of the three average readings of the technical duplicates. B. Heme influx in heme-starved cells. *hem1*^*-/-*^ cells (KC1088) expressing the three HS1 heme sensors were grown overnight in YPD + ferrozine in the absence of ALA. Cells ceased growth because of heme deficiency at an OD_600_ density of 1-3. The cultures were then diluted to a density of 1, and the indicated amounts of hemin or hemoglobin were added. At the indicated time points, a sample from each culture was measured as in A. Note that the time-scales are not identical. C. Heme influx into *hem1*^*-/-*^ cells lacking the main hemophore Pga7. *hem1*^*-/-*^ *pga7*^*-/-*^ cells (KC1111) expressing the three HS1 heme sensors were depleted of heme as in A and exposed to 10 µM heme or 5 µM hemoglobin for the indicated amounts of time. Sample measurement was as described for 3A. D. Effect of sodium azide on heme influx into *hem1*^*-/-*^ cells lacking the main hemophore Pga7. *hem1*^*-/-*^ *pga7*^*-/-*^ cells (KC1111) were depleted of heme as in Fig. 3A and exposed to 10 µM heme for the indicated amounts of time, in the presence or absence of 20 mM sodium azide as indicated. The graph depicts WT and M7A fluorescence ratios normalized to DM fluorescence ratio. The primary data are shown in Fig. S5.

In the next set of experiments, we monitored the kinetics of influx of LH into the cytoplasm in a strain completely dependent upon an external heme supply. The *hem1*^*-/-*^ mutant was grown overnight in the absence of ALA to deplete it of heme. The cells were then resupplied with either hemin or hemoglobin, and the heme sensor fluorescence was measured at different time points. As shown in Fig. 3B, the initial fluorescence ratio of the three sensor variants was similar, indicating ∼0 % heme occupancy of all sensors as expected, since these cells arrested growth due to heme depletion. Addition of 2.5 µM hemin led to an increase in WT HS1 heme occupancy from 0% to ∼50% after 45 min, and by 135 min the sensor was quantitatively saturated with heme (> 95% heme loaded). Between 135 min and 225 min, exogenous heme even began to populate the lower affinity HS1-M7A sensor, leading to a heme occupancy of 37% after 225 min. In stark contrast, upon hemoglobin addition (2.5 µM, *i*.*e*. 10 µM of heme equivalents), WT HS1 heme occupancy went from 0% to ∼40% only after 150 min, and was never quantitatively saturated with heme, reaching ∼85% heme occupancy only after 300 min. Moreover, heme from hemoglobin never contributed towards the loading of HS1-M7A (Fig. 3B). In parallel, we measured resumption of growth in these cultures. The cultures exposed to hemoglobin did not increase in density even after 300 min, whereas the cultures exposed to hemin resumed growth at 150-180 min (Fig. S4). We conclude that hemin is taken up much faster into the cells than hemoglobin heme. This could be due to the presence of an alternative pathway for free hemin, or it could reflect the fact that before entering the hemophore transfer cascade, hemoglobin heme must first be extracted from the globin in a rate determining step.

The notion that hemin can be taken up by a pathway different from the CFEM hemophores pathway is supported by the observation that at higher concentration, hemin can serve as an iron source even in the absence of *PGA7*, a gene that is essential for hemoglobin-iron utilization (Kuznets *et al*., 2014; Pinsky *et al*., 2020). To analyze the characteristics of this alternative pathway, we measured influx of heme into a double *hem1*^*-/-*^ *pga7*^*-/-*^ mutant depleted of endogenous heme, using either 10 µM hemin or 5 µM hemoglobin as heme sources (Fig. 3C). As expected, hemoglobin did not serve as source of heme influx into the *pga7*^*-/-*^ strain even after 4 hours. Hemin, in contrast, did cause population of the high-affinity HS1 sensor within 1 hour, however the medium-affinity M7A sensor remained unbound even after 3 hours, in spite of the high external hemin concentration. We conclude that while an alternative pathway exists for uptake of free hemin into the cell, it is relatively slow and cannot account for the rapid uptake of hemin into the *PGA7*^*+/+*^ cells. Thus, the CFEM hemophore cascade is probably responsible for the rapid heme uptake detected in Fig. 3B.

We also tested whether the non-Pga7 dependent heme influx is energy-dependent by assaying the effect of sodium azide, a mitochondrial uncoupler that causes rapid depletion of cellular energy stores, on hemin uptake in the *pga7*^*-/-*^ cells. However, we found that the non-binding DM HS1 construct showed reduced fluorescence ratios in the presence of azide, indicating that azide reduces sensor fluorescence unrelated to heme binding (Fig. S5). In order to account for this reduction caused by azide, we calculated the results of HS1 and M7A fluorescence ratios normalized to DM fluorescence ratio (Fig. 3D). The results indicate that in the absence of Pga7, the addition of azide prevented heme influx in the first hour of exposure, whereas after 3 hours, about half of the HS1 sensor was bound by heme (Fig. 3D). This suggests that the Pga7-independent heme uptake pathway is energy dependent. The limited heme influx detected after 3 hours could be due to residual activity of this pathway, or to passive diffusion of heme into the cell.

### Role of Hmx1 in cellular heme levels

Since we found that the heme oxygenase Hmx1 had only a minor effect on heme-iron utilization, we tested its effects on intracellular heme levels in the absence or presence of different external heme concentrations. All cultures were grown in the presence of 1 mM ferrozine, to impose partial iron limitation in order to activate the heme uptake system, and the cultures were then exposed for 5 to 7 hours to different concentrations of hemin or hemoglobin. In the absence of added heme, we found that HS1 had similar heme occupancy (∼30%) in the WT, *hmx1*^*-/-*^, and *hmx1*^*-/-*^ <*HMX1*> reintegrant cells (Fig. 4A). Addition of hemin led to significantly elevated intracellular heme in *hmx1*^*-/-*^ cells as compared to wild-type and the *hmx1*^*-/-*^ <*HMX1*> reintegrant cells (Fig. 4A). For instance, supplementation with 5 µM hemin resulted in the near saturation of the lower affinity HS1-M7A sensor in cells lacking Hmx1. The same experiment done with a different *HMX1* wild-type and mutant strain set gave a very similar result (Fig. S6). We thus conclude that in the absence of *HMX1*, exposure to hemin in the medium leads to significantly higher levels of intracellular heme.

**FIG. 4.**
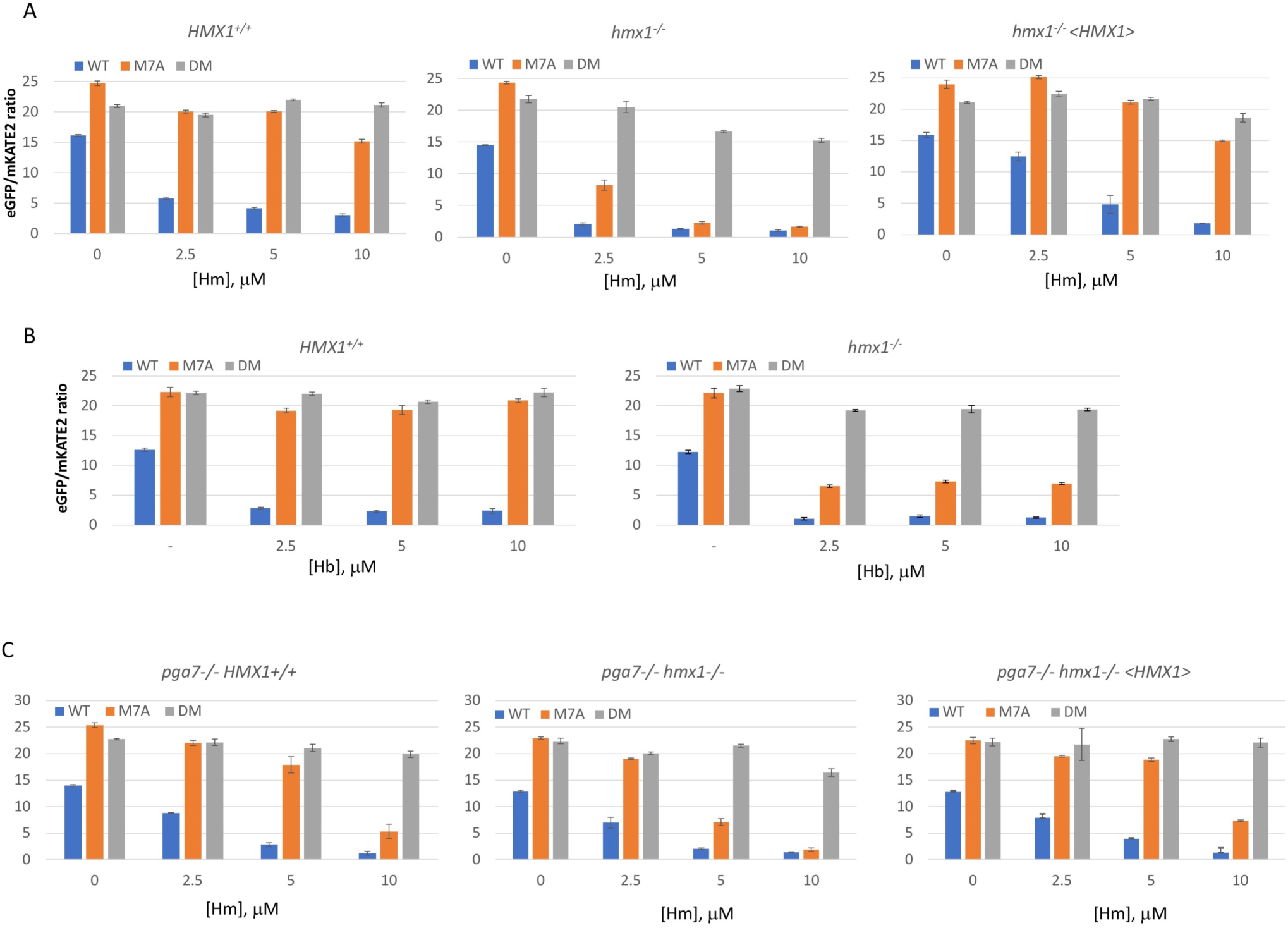
Cytoplasmic heme levels are increased in the *hmx1*^*-/-*^ mutant exposed to hemin or hemoglobin in the medium. A. *HMX1*^*+/+*^ (KC1262), *hmx1*^*-/-*^ (KC1251) or *hmx1*^*-/-*^ <*HMX1*> reintegrant cells (KC1267) were grown in YPD + 1 mM ferrozine and the indicated hemin concentrations for 5 hours. B. *HMX1*^*+/+*^ (KC1262) or *hmx1*^*-/-*^ (KC1251) cells were grown in YPD + 1 mM ferrozine and the indicated hemoglobin concentrations for 5 hours. C. *pga7*^*-/-*^ (KC1276), *pga7*^*-/-*^ *hmx1*^*-/-*^ (KC1253) or *pga7*^*-/-*^ *hmx1*^*-/-*^ *<HMX1>* reintegrant cells (KC1281) were exposed to the indicated external hemin concentrations for 5 hours. All samples were measured as in 3A.

We next tested the effect of hemoglobin in the medium on heme homeostasis in the *HMX1* wild-type and mutant strains. In the case of hemoglobin, we found that concentrations from 2.5 µM to 10 µM gave the same effect on intracellular heme levels; the wild-type HS1 sensor remained nearly quantitatively saturated in both WT and *hmx1*^*-/-*^ cells, whereas the heme occupancy of the HS1-M7A sensor went from < ∼15% to ∼60% upon deletion of *HMX1* (Fig. 4B). Thus, while the *hmx1*^*-/-*^ mutant still has higher levels of cytoplasmic heme in the presence of hemoglobin in the medium, those levels, as well as heme levels in the wild-type cells, are independent of the external hemoglobin concentration in the medium. This suggests that the hemoglobin heme import pathway is already working at maximum velocity in 2.5 µM hemoglobin, but not in 2.5 µM hemin. This implies that the rate-limiting step in the hemoglobin-heme import pathway is the extraction of heme from hemoglobin by the CFEM hemophore cascade.

Unlike hemoglobin, which can enter the cells via the CFEM hemophore pathway only, hemin can penetrate the cell via one or more additional alternative pathways (Kuznets *et al*., 2014; Pinsky *et al*., 2020) (Fig. 3C). To test hemin homeostasis when the CFEM heme uptake pathway is disrupted, we deleted *HMX1* in the *pga7*^*-/-*^ mutant background, lacking the main CFEM hemophore Pga7. In the *pga7*^*-/-*^ *HMX1* wild-type and reintegrant strains, increasing external hemin causes a dose-dependent increase in heme loading of the HS1 sensor and substantial loading of the HS1-M7A sensor at the highest hemin concentration (10 µM) (Fig. 4C). Levels of heme bound were however lower than in the *PGA7* wild-type and reintegrant cells at equivalent concentrations (compare to Fig. 4A). In the *pga7*^*-/-*^ *hmx1*^*-/-*^ strain, at equivalent external heme concentrations, both the HS1 and HS1-M7A heme sensors were more populated with heme than in the *HMX1* wild-type and reintegrant strains, indicating that absence of Hmx1 caused higher accumulation of heme in the cytoplasm in the absence of Pga7 as well.

### Hmx1 protects cells against extracellular heme toxicity

Since increased free heme concentrations can be toxic to cells, and since we found that more hemin penetrates into the cells with increasing external concentrations, we next tested whether the *hmx1*^*-/-*^ strain is more sensitive to heme in the medium. As can be seen in Fig. 5, a limited growth inhibition was detected in 5 µM hemoglobin in the *hmx1*^*-/-*^ strain compared to the wild-type and reintegrant strains, and an increasing inhibition of growth was detected when going from 30 µM to 50 µM and 100 µM of hemin. This supports the conclusion that a major role of Hmx1 is to prevent cytoplasmic heme from reaching toxic levels when the cells are utilizing external heme.

**FIG. 5.**
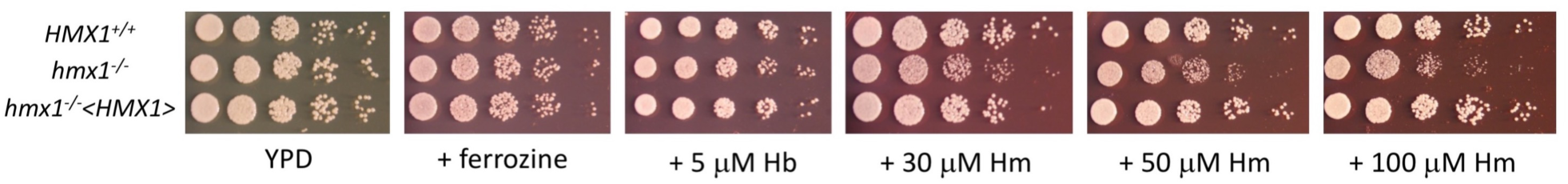
The *hmx1*^*-/-*^ mutant is sensitive to excess heme in the medium. Dilutions of *HMX1*^*+/+*^ strain (KC1065), of a *hmx1*^*-/-*^ mutant (KC1066), and of a *hmx1*^*-/-*^ <*HMX1*> reintegrant (KC1067) were spotted on YPD + 1 mM ferrozine plates containing the indicated amounts of hemoglobin or hemin, and incubated at 30°C for 1 day.

## DISCUSSION

While *C. albicans* had been known to utilize heme as an iron source, we have shown here that *C. albicans* can utilize heme as a cellular heme source as well. In fungi, a pathway for utilization of external heme as a heme source had been previously described only in the fission yeast *S. pombe* (Mourer *et al*., 2015; Labbé *et al*., 2020). It is not clear yet whether *S. pombe* can also use heme as an iron source: the lack of a heme oxygenase gene homolog in *S. pombe* has been cited to support the assumption that this organism cannot utilize heme as an iron source (Labbé *et al*., 2020), but the limited role of heme oxygenase that we detected in heme-iron utilization in *C. albicans* undermines this reasoning. What is clear however is that *S. pombe*, a heme prototroph, has a dedicated pathway for heme uptake, and that this pathway is induced under iron starvation conditions (Mourer *et al*., 2015). Even if not used as an iron source, uptake of external heme under iron scarcity conditions could help the *S. pombe* cells save the iron that would otherwise be needed for heme biosynthesis.

In *C. albicans*, the heme uptake pathway is strongly induced under iron starvation conditions (Weissman and Kornitzer, 2004; Chen *et al*., 2011; Kuznets *et al*., 2014; Nasser *et al*., 2016; Roy and Kornitzer, 2019). While heme can in fact be used as an alternative iron source by *C. albicans*, the ability shown here to use external heme as a heme source would save the cells the need for the elemental iron required in heme synthesis, while in addition also saving the metabolic cost of protoporphyrin synthesis. This raises the question whether, even in the presence of iron, *C. albicans* cells could utilize external heme. One striking observation is that, in addition to iron starvation conditions, the CFEM hemophores involved in heme uptake are also among the most strongly induced genes under hyphal induction conditions (Azadmanesh *et al*., 2017; Roy and Kornitzer, 2019). *C. albicans* is a dimorphic pathogen, capable of switching from a yeast to a hyphal (mold) morphology under a variety of conditions such as elevated temperature, the presence of serum, the bacterial and fungal cell wall precursor N-acetyl glucosamine, among others (Sudbery *et al*., 2004; Basso *et al*., 2019; Kornitzer, 2019). While there is no evidence that the CFEM hemophores are required for hyphal morphogenesis, it is possible that hyphal-inducing conditions signify a heme-rich environment, be it the gut lumen or the mucosal surface, which would promote heme uptake even when iron is relatively abundant.

The ability of *C. albicans* to take up external heme and utilize it as either a cellular heme source or as an iron source, led us to question what the additional requirements are for utilization of heme as an iron source. Release of the iron from heme requires breakdown of the protoporphyrin IX ring. The main heme degrading activity in eukaryotic cells is heme oxygenase (Maines, 1988; Ponka, 1999). A heme oxygenase homolog was detected in both *S. cerevisiae* and in *C. albicans*, and was shown to carry heme-degrading activity (Protchenko and Philpott, 2003; Santos *et al*., 2003; Pendrak *et al*., 2004). Mammalian heme oxygenases are anchored to the cytoplasmic face of the ER membrane (Yoshida and Sato, 1989). For *S. cerevisiae* Hmx1, this localization is supported by immunolocalization to the ER and by the observation that the heme turnover defect in the *hmx1Δ* strain is complemented by expression in the cytoplasm of a bacterial heme oxygenase, *Corynebacterium diphtheriae* HmuO (Protchenko and Philpott, 2003). In *C. albicans*, the CFEM pathway of heme import probably channels the heme to the vacuole, and *S. cerevisiae* cells expressing the CFEM protein Rbt51/Pga10 and therefore able to utilize hemoglobin as iron source, require the vacuolar iron transporter Smf3, suggesting that iron is released from the heme in the vacuole (Weissman *et al*., 2008). Assuming that the heme uptake pathway and Hmx1 localization are similar in *C. albicans*, Hmx1 was therefore not expected to play a major role in heme-iron utilization. This is in fact what we found: a *hmx1*^*-/-*^ mutant is only weakly defective for both hemoglobin and hemin utilization. This is in apparent contradiction to previously published results, which found a significant defect of a *C. albicans hmx1*^*-/-*^ mutant in growth on hemin as an iron source (Santos *et al*., 2003). However, the hemin concentration used in that report, at 50 µM, was quite high, and we found that at such concentrations, hemin starts to become toxic in a *hmx1*^*-/-*^ mutant (Fig. 5). It is thus possible that the previous report mainly detected an effect of Hmx1 on detoxification of excess heme rather than on heme-iron utilization.

Although Hmx1 is not necessary for heme-iron acquisition, we do find that like in other eukaryotes, *C. albicans* Hmx1 is involved in heme homeostasis. Under conditions of partial iron limitation (1 mM ferrozine), but in the absence of external hemin, we did not detect any effect of *HMX1* deletion on cytoplasmic LH levels. The function of Hmx1 became apparent when heme was provided in the medium as hemoglobin or as hemin: the *hmx1*^*-/-*^ mutant accumulated higher cytoplasmic LH levels than the wild-type strain. This accumulation probably accounts for the selective heme sensitivity of the *hmx1*^*-/-*^ mutant.

Thus, our results indicate that Hmx1 is specifically involved in maintaining heme homeostasis in the presence of excess extracellular heme, but it is not essential for utilization of the iron from external heme.

The results presented here raise several new questions. For one, the non-essentiality of Hmx1 in heme-iron utilization implies that fungal cells must carry an additional heme-degrading activity, possibly in the vacuole. This activity remains to be characterized.

Secondly, to the extent that import of heme from hemoglobin into the cell involves endocytosis into the vacuole (Weissman *et al*., 2008), the appearance of external heme in the cytoplasm suggests the existence of a yet-uncharacterized vacuolar transporter, similar perhaps to the Abc3 transporter in *S. pombe* (Mourer *et al*., 2017), that exports heme from the vacuole to the cytoplasm.

In summary, the use of novel genetically encoded heme sensors that enable monitoring of cytoplasmic LH concentrations under different conditions allowed us to generate a more detailed picture of the mechanisms and kinetics of external heme uptake in *C. albicans*. We confirmed that hemoglobin heme is exclusively imported via the CFEM hemophore cascade, and found that heme extraction from hemoglobin is probably the rate-limiting step of this pathway. Hemin import into the cell is more rapid, both because it bypasses the heme extraction step, and because hemin can enter the cell via an alternative, as yet uncharacterized, energy-dependent pathway. External heme, whether from hemoglobin or hemin, is channeled to the cytoplasm, and we find that the main role of the heme oxygenase Hmx1 is to prevent heme toxicity in the face of this external heme influx. The different functions involved in import and homeostasis of heme in *C. albicans* underscore its multiple roles as essential protein cofactor, possible iron source, and potentially toxic molecule.

## MATERIALS AND METHODS

### Media

Cells were grown in YPD medium (1% yeast extract, 2% bacto-peptone, 2% glucose, tryptophan 150 mg/l) supplemented with the ion chelators ferrozine or bathophenanthroline sulfonate (Sigma) at 1 mM, and hemin from a 2 mM stock in 50 mM NaOH, or hemoglobin from a 0.5 mM stock in PBS (Phosphate-buffered saline: 137 mM NaCl, 2.7 mM KCl, 10 mM Na_2_HPO_4_, 2 mM KH_2_PO_4_, pH = 7.4). Hemin was obtained from Frontier Scientific, and bovine hemoglobin (H2500) and HSA (A3782) from Sigma. Synthetic Complete medium contains, per liter, Yeast Nitrogen Base (USBiological) 1.7 g, (NH_4_)_2_SO_4_ 5 g, the 20 amino acid plus adenine and uridine, 0.1g each except leucine, 0.2 g, glucose 20 g, and 0.2 mM inositol.

### Strains

*C. albicans* strains are listed in Table 1. KC947, KC965 contain the CaCas9-expressing plasmid pV1025 (Vyas *et al*., 2015) at the *ENO1* locus. In KC 1068 and KC1069, the two *HMX1* alleles were mutagenized by introduction of a CRISPR-mediated G74 →CA mutation, introducing an SphI site and a frameshift, using KB2587 as gRNA-expressing plasmid. In KC1088 and KC1111, the two *HEM1* alleles were mutagenized by introduction of a CRISPR-mediated ΔC197, A199T mutation, introducing a SacI site and a frameshift, using KB2588 as gRNA-expressing plasmid. KC1243 was constructed by sequential replacement of the *PGA7* allele with hisG using the KB2025 “URA3 blaster” plasmid (Kuznets *et al*., 2014). KC1251, KC1253 were generated by simultaneous replacement of both *HMX1* alleles with CdLEU2 using a transient CRISPR-promoted homologous recombination method (Min *et al*., 2016) as follows: fusion PCR was used to obtain a PCR fragment with the CdLEU2 marker flanked on each side by 500 bp of *HMX1* flanking sequences, as described in (Noble and Johnson, 2005). This fragment was then co-transformed with a CaCas9-expressing PCR fragment and a *HMX1* gRNA-expressing PCR fragment amplified from pV1025 and KB2587, respectively. KC1262, KC1276 were made Leu+ by transformation with a *LEU2* PCR fragment. KC1267, KC1281 have a *HMX1* wild-type allele reintroduced at the *HMX1* locus by transformation with plasmid KB2721 digested with SpeI.

**Table 1.** List of *Candida albicans* strains.

| Name | Genotype | Origin |
| --- | --- | --- |
| KC2 = CAI4 | *ura3Δ::imm434/ura3Δ::imm434* | (Fonzi and Irwin, 1993) |
| KC68 | KC2 *ccc2Δ::hisG/ccc2Δ::hisG* | (Weissman *et al.*, 2002) |
| KC274 = SN148 | *ura3Δ::imm434/"" iro1Δ::imm434/""*<br>*his1Δ/"" leu2Δ/"" arg4Δ/""* | (Noble and Johnson, 2005) |
| CaSS1 | CAI4 *his3Δ/his3Δ LEU2/leu2::tetR-*<br>*GAL4AD* | (Roemer *et al.*, 2003) |
| TEToff-HEM1 | CaSS1 *hem1::HIS3/TETpr::HEM1*<br>*NAT<sup>r</sup>* | (Roemer *et al.*, 2003) |
| KC482 | KC2 *pga7Δ/pga7Δ* | (Kuznets *et al.*, 2014) |
| KC1065 = SC5314 |  | (Navarathna and Roberts, 2010) |
| KC1066 = DLR2 | *hmx1Δ/hmx1Δ* | (Navarathna and Roberts, 2010) |
| KC1067 = DLR3 | *hmx1Δ/hmx1Δ <HMX1>* | (Navarathna and Roberts, 2010) |
| KC947 | KC68 *ENO1/ENO1::CaCAS9* | This work |
| KC965 | KC2 *ENO1/ENO1::CaCAS9* | This work |
| KC1068 | KC947 *hmx1<sup>-</sup>/hmx1<sup>-</sup> NAT<sup>r</sup>* | This work |
| KC1069 | KC965 *hmx1<sup>-</sup>/hmx1<sup>-</sup> NAT<sup>r</sup>* | This work |
| KC1088 | KC965 *hem1<sup>-</sup>/hem1<sup>-</sup> NAT<sup>r</sup>* | This work |
| KC1111 | KC482 *hem1<sup>-</sup>/hem1<sup>-</sup>*<br>*ENO1/ENO1::CaCAS9-NAT<sup>r</sup>* | This work |
| KC1251 | KC274 *hmx1*Δ::LEU2/ *hmx1*Δ::LEU2 | This work |
| KC1262 | KC274 *leu2*Δ/LEU2 | This work |
| KC1267 | KC1251 *hmx1*Δ::LEU2/HMX1 NAT<sup>r</sup> | This work |
| KC1243 | KC274 *pga7*Δ/ *pga7*Δ | This work |
| KC1276 | KC1243 *leu2*Δ/LEU2 | This work |
| KC1253 | KC1243 *hmx1*Δ::LEU2/ *hmx1*Δ::LEU2 | This work |
| KC1281 | KC1253 *hmx1*Δ::LEU2/HMX1 NAT<sup>r</sup> | This work |

### Plasmids

KB2587 expresses *HMX1* sgRNA sequence ATACCTGCTAAAACTGACGT (52-63) from plasmid pV1090 (Vyas *et al*., 2015). KB2588 expresses *HEM1* sgRNA sequence TGTAGAAGGAGTAGCAGCGG (200-219) from plasmid pV1090. KB2721 contains a *HMX1* genomic fragment extending from -1550 to + 959, ApaI to NotI, into KB2248. KB2248 is pBSII KS+ containing a NAT1 SacI-NotI PCR fragment amplified from pJK795 (Shen *et al*., 2005).

CaHS1 sensor plasmids: in a first stage, we made a synthetic HS1 gene (Hanna *et al*., 2016) modified for expression in *C. albicans* by substituting all 21 CTG codons (Genescript), called CaHS1. We next tested expression of this gene under five promoters, *ACT1p, TPI1p, ENO1p, MAL2p, CUP1p* under different conditions. The *ENO1* promoter gave the most consistent CaHS1 fluorescence and was used in all assays shown. KB2636 consists of the vector BES116 (Feng *et al*., 1999) with the *ENO1* promoter region extending from -955 to -1 between NotI-SpeI, and CaHS1 cloned between SpeI-ApaI. KB2669 is KB2636 with the M7A mutation introduced by PCR-mediated site-directed mutagenesis. KB2671 is KB2669 with an added H102A mutation, i.e. the double mutant. All three plasmids were transformed in the appropriate strains by integration at the *ADE2* locus.

### Growth assays

Liquid assays: Overnight cultures grown in YPD were diluted into a series of two-fold dilutions of hemin or hemoglobin, in YPD + ferrozine or BPS. Cells were inoculated in flat-bottomed 96-well plates at OD_600_=0.00001, 150 µl per well. Plates were incubated at 30°C on an orbital shaker at 60 rpm and growth was measured by optical density (OD_600_) after 2 and 3 days with an ELISA reader. Cells were resuspended with a multi-pipettor before each reading. Each culture was done in triplicate.

Spot assays: Overnight cultures grown in YPD were serially diluted in PBS. The culture was first diluted 1:100, and then sequentially 1:5 five times. 3 µl of each dilution was spotted on the indicated agar plates.

### Heme sensor binding assays

Each strain to be tested was transformed with the sensor plasmids KB2636 (WT HS1), KB2669 (M7A), KB2671 (DM), as well as a vector plasmid. Three cultures each were set up for each sensor plasmid, and a single vector plasmid culture. Overnight cultures were diluted to O.D. _600_ = 0.2, and left to grow to O.D. _600_ ∼ 1, but not over 2 (5-7 h), before taking measurements. *hem1*^*-/-*^ strains grown without ALA were diluted to O.D. _600_ = 0.8, refed with the indicated heme sources, and measurements were taken at the indicated times. For fluorescence measurements, cells were pelleted, washed once with PBS, resuspended to O.D. _600_ = 5, and 0.2 ml were placed in duplicate in a black 96 well flat bottom plates (Nunc or Grenier Fluorotrac). Fluorescence was measured with a Tecan infinite 200 Pro machine, with eGFP: ex. 480 nm (9 nm bandwidth), em. 520 nm (20 nm bandwidth), and mKATE2 ex. 588 nm (9 nm bandwidth), em. 620 nm (20 nm bandwidth). For Fig. S3B only: fluorescence was measured using a Biotek Synergy MX plate with eGFP: ex. 480 nm (9 nm bandwidth), em. 520 nm (9 nm bandwidth), and mKATE2 ex. 588 nm (9 nm bandwidth), and em. 620 nm (9 nm bandwidth). For each data point, each culture was measured twice (technical duplicate), the vector-only culture reading was substracted from all other readings, and the ratio of eGFP to mKATE2 was calculated. Shown are mean and standard deviation of 3 biological replicates.

### Fractional heme sensor saturation measurement

In order to relate the sensor fluorescence ratio to the fractional heme saturation of the sensor for quantitative LH measurements, the following equation is used

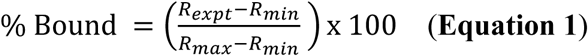

The amount of heme bound to the sensor, % Bound, can be quantified by determining the sensor eGFP/mKATE2 fluorescence ratio under any given experimental condition, *R*_expt_, relative to the eGFP/mKATE2 fluorescence ratios when the sensor is 0% (*R*_min_) and 100% (*R*_max_) bound to heme, respectively. *R*_max_ is assumed to be close to 0, based on the > 99% efficiency of energy transfer between GFP and heme. This was empirically confirmed in cells having high cytoplasmic LH concentrations such as the *hmx1*^*-/-*^ mutant. *R*_min_ can be determined by growing a parallel culture of heme deficient *hem1*Δ cells expressing the heme sensor or from cells expressing HS1-M7A,H102A, which cannot bind heme.

### Characterization of labile heme sensors by microscopy

*C. albicans* expressing CaHS1 plasmids KB2636 (CaHS1), KB2669 (HS1-M7A), KB2671 (HS1-M7A,H102A = DM) and a vector-only plasmid were grown overnight in SC media at 37 °C. Cells were washed in PBS and resuspended at an O.D. _600_ = 0.7 in PBS. Cells were immediately loaded onto a CellASIC Onix Y04C yeast microfluidics plate. Cells were imaged on a Zeiss LSM 710 NLO confocal microscope with a 40X 1.3 numerical aperture oil-immersion objective. A 488 nm laser was used to excite GFP and a 561 nm laser was used to excite mKATE2. Emission filters for GFP and mKATE2 were 493 – 536 nm and 566 – 685 nm respectively. *C. albicans* expressing the vector only plasmid was used for background subtraction. Images were collected with Zeiss software and analyzed with ImageJ. (Rasband, W.S., ImageJ, U. S. National Institutes of Health, Bethesda, Maryland, USA, https://imagej.nih.gov/ij/, 1997-2018.)

## ACKNOWLEDGMENTS

This research was supported by a US-Israel Binational Science Foundation grant #2017228 (to AR and DK), a US National Institutes of Health grant ES025661 (to AR), a US National Science Foundation grant 1552791 (to AR), and an Israel Science Foundation grant 487/15 (to DK). We thank David Roberts, Valmik Vyas, Julia Koehler and Andre Nantel for strains and plasmids. We wish to acknowledge the core facilities at the Parker H. Petit Institute for Bioengineering and Bioscience at the Georgia Institute of Technology for the use of their shared equipment, services and expertise.

## AUTHOR CONTRIBUTIONS

Conceived and designed the experiments: DK, ARR. Performed the experiments: ZW, MP, RKD. Analysed the data: ZW, MP, RKD, ARR, DK. Contributed reagents/materials/analysis tools: RKD, ARR. Supervised the project: ARR, DK. Drafted the manuscript: DK. All authors critically read and revised the manuscript.

## SUPPLEMENTARY FIGURES

**FIG. S1.**
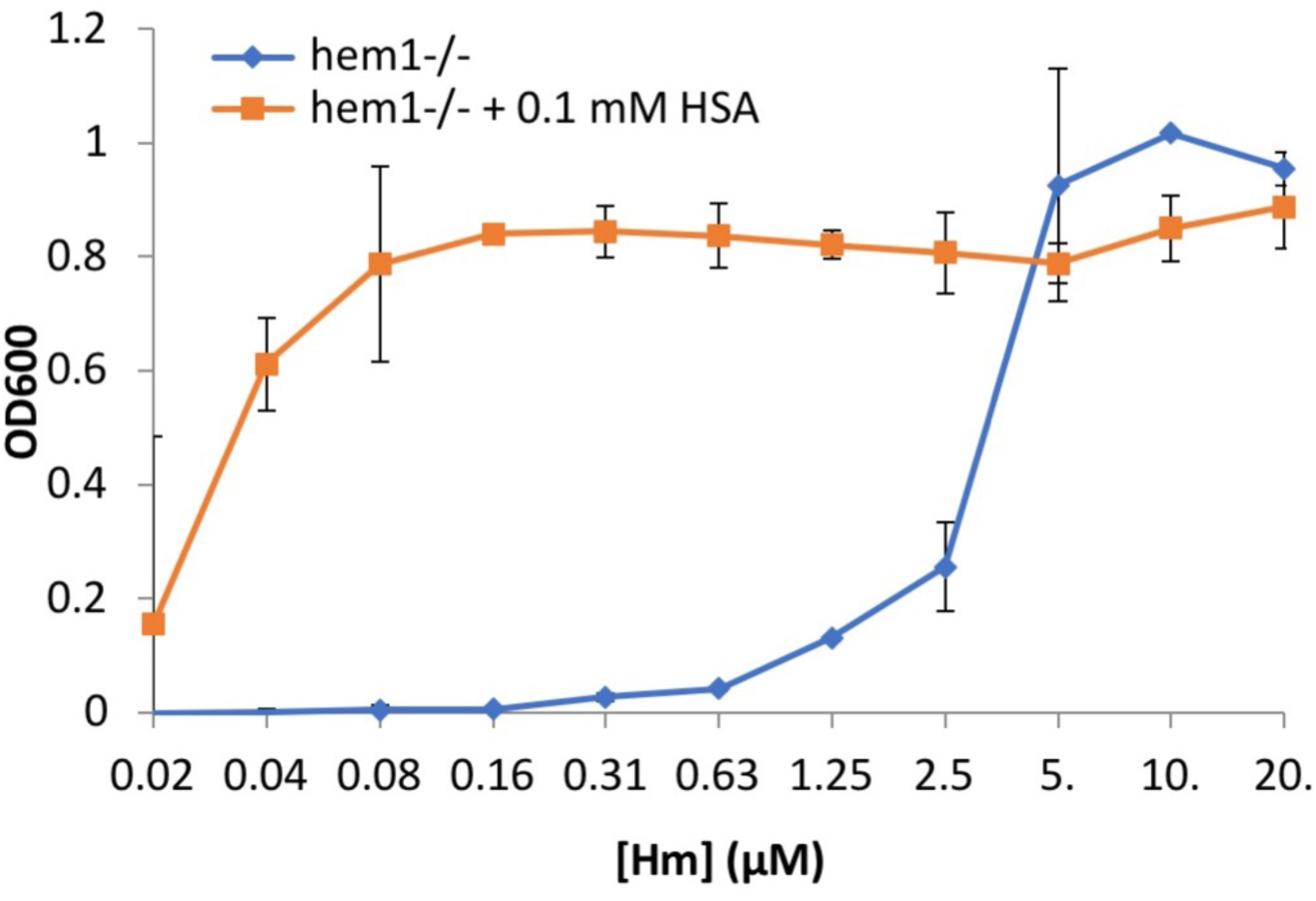
*C. albicans* can efficiently utilize albumin-complexed heme as heme source. The *hem1*^*-/-*^ strain (KC1088) was grown in YPD medium + 1 mM ferrozine, with or without addition of 0.1 mM human serum albumin (HSA). The cultures were grown at 30°C for 3 days. Each data point represents the average of three independent cultures, and the error bars represent the standard deviation.

**FIG. S2.**
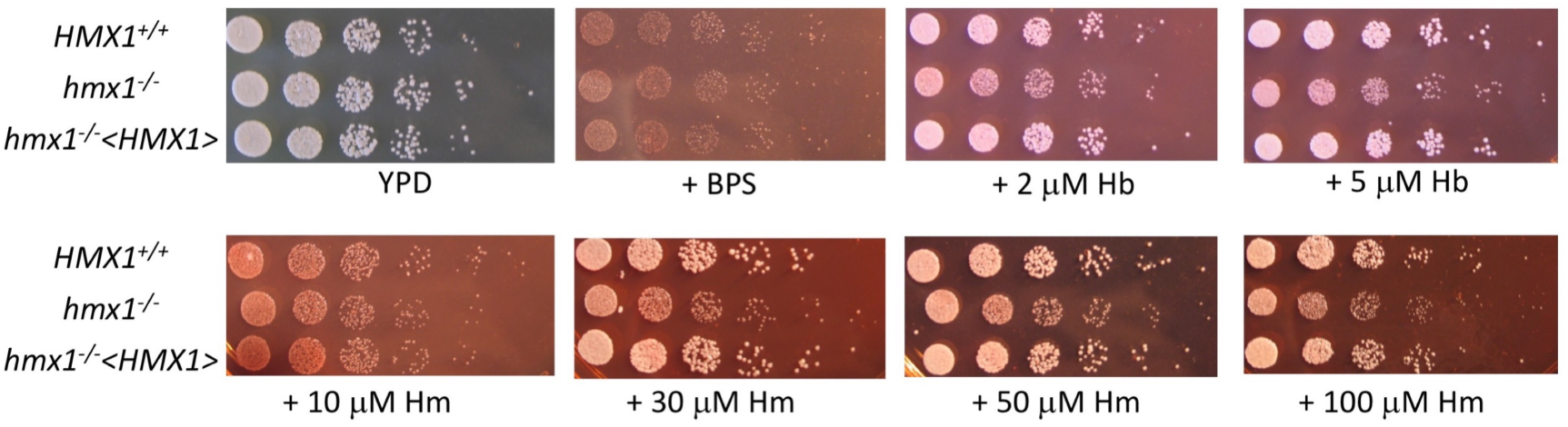
Dilutions of a wild-type strain (KC274), of a *hmx1*^*-/-*^ mutant (KC1251), and of a *hmx1*^*-/-*^ <*HMX1*> reintegrant (KC1267) were spotted on YPD or on YPD + 1 mM BPS, without or with the additions of the indicated hemoglobin and hemin concentrations, and incubated at 30°C for 2 days.

**FIG. S3.**
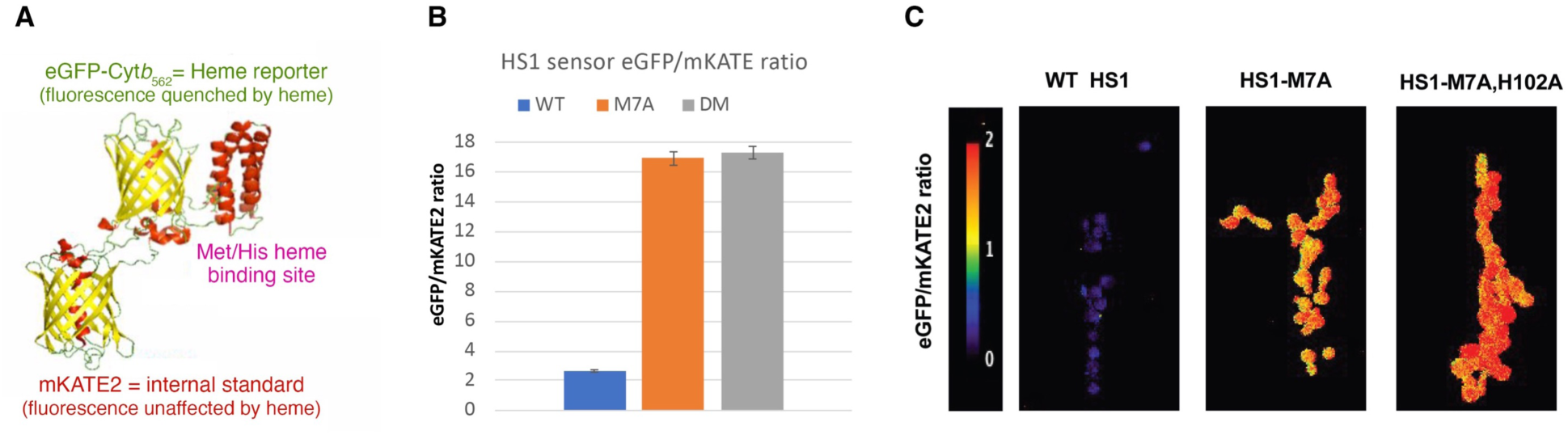
The HS1 heme sensor. A. Schematic depiction of the HS1 heme sensor protein structure. B. *C. albicans* cells expressing codon-adapted HS1 genes on plasmids KB2636 (wild-type HS1), KB2669 (HS1-M7A), KB2671 (HS1-M7A,H102A=DM) or vector-only were grown overnight in SC medium supplemented with ergosterol and Tween 80. The cells were then diluted in the same medium and grown to an O.D. between 1 and 2. Fluorescence was measured as described in Methods. The graph depicts the ratio between the eGFP fluorescence and the mKATE2 fluorescence. C. The same strains as in B were grown overnight in SC medium at 37°, then visualized by confocal microscopy as described in Methods.

**FIG. S4.**
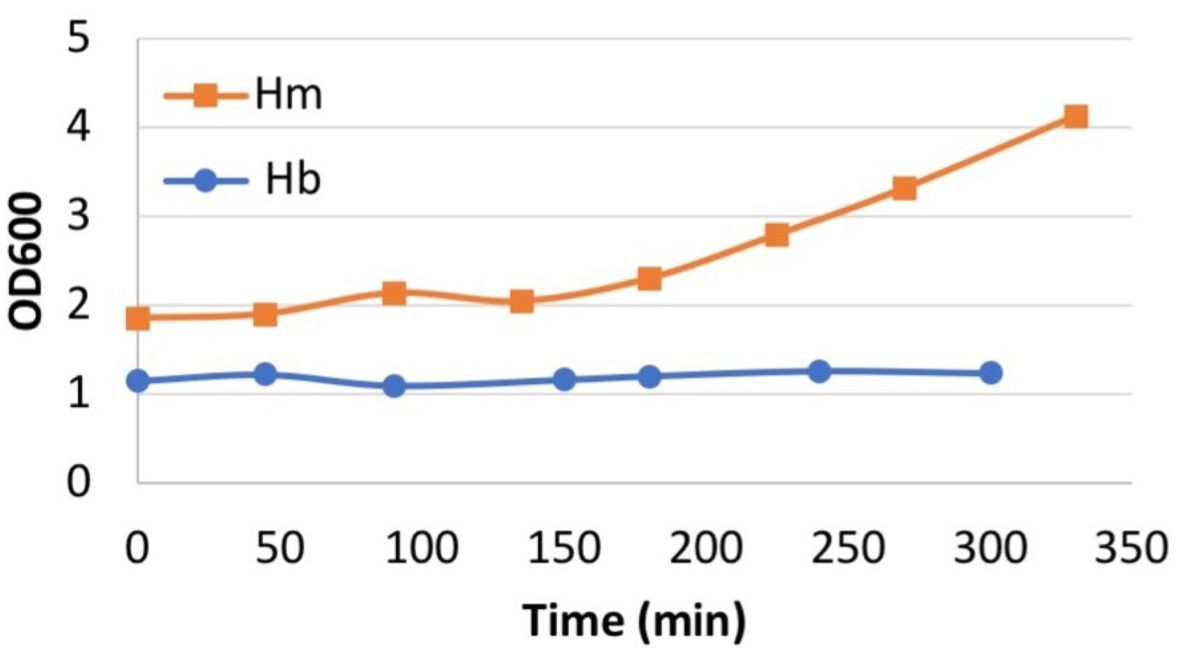
Growth recovery after addition of hemin or hemoglobin to heme-starved *hem1*^*-/-*^ cells. Density of cultures starved for heme and resupplied with hemin or hemoglobin. The cultures analyzed in Fig. 3B were measured for optical density after addition of hemin or hemoglobin. Each data point is the average of ten cultures.

**FIG. S5.**
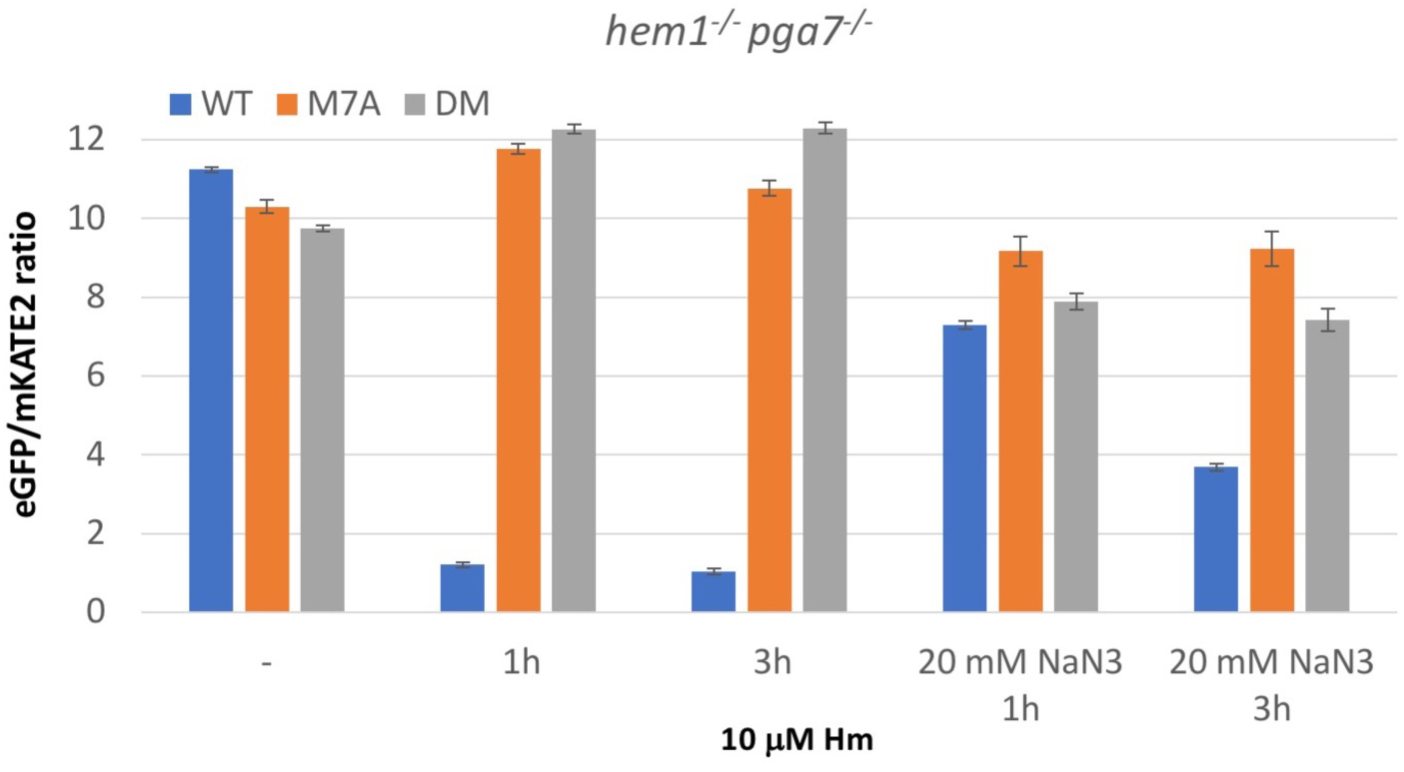
Heme influx into *hem1*^*-/-*^ cells lacking the main hemophore Pga7. *hem1*^*-/-*^ *pga7*^*-/-*^ cells (KC1111) were depleted of heme as in Fig. 3A and exposed to 10 µM heme for the indicated amounts of time, in the presence or absence of 20 mM sodium azide as indicated. Sample measurement was as described for 3A.

**FIG. S6.**
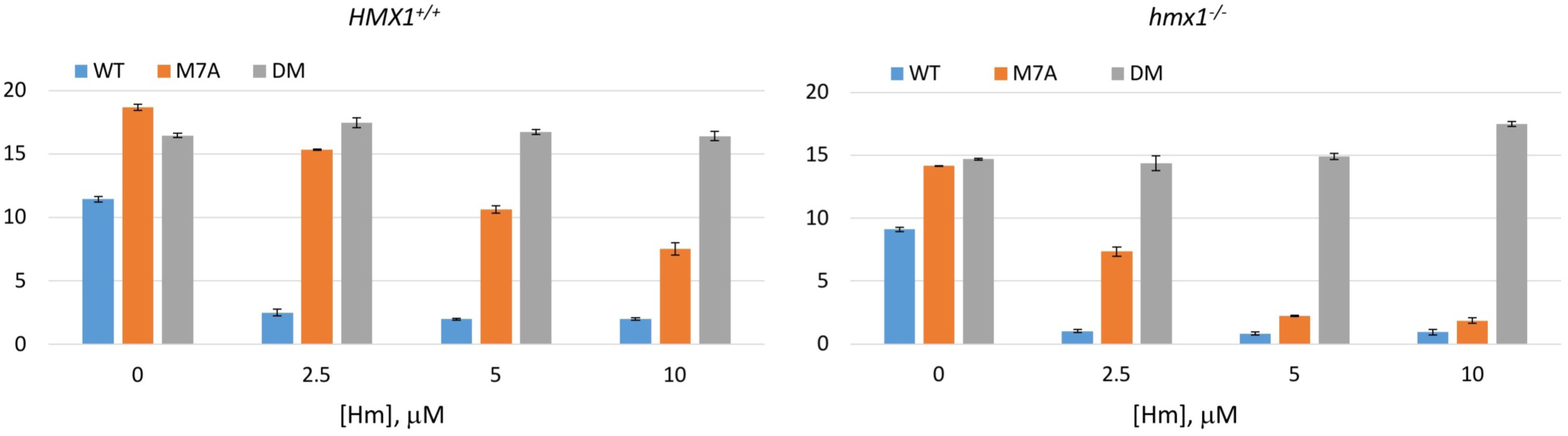
Cytoplasmic heme levels are increased in a *hmx1* mutant exposed to extracellular hemin. *HMX1*^*+/+*^ (KC965) and *hmx1*^*-/-*^ (KC1069) cells were grown 7 h in YPD + 1 mM ferrozine and the indicated amount of hemin.

